# Feasibility of Using an Armband Optical Heart Rate Sensor in Naturalistic Environment

**DOI:** 10.1101/2022.10.03.510715

**Authors:** Hang Yu, Michael Kotlyar, Sheena Dufresne, Paul Thuras, Serguei Pakhomov

## Abstract

Consumer-grade heart rate (HR) sensors including chest straps, wrist-worn watches and rings have become very popular in recent years for tracking individual physiological state, training for sports and even measuring stress levels and emotional changes. While the majority of these consumer sensors are not medical devices, they can still offer insights for consumers and researchers if used correctly taking into account their limitations. Multiple previous studies have been done using a large variety of consumer sensors including Polar^®^ devices, Apple^®^ watches, and Fitbit^®^ wrist bands. The vast majority of prior studies have been done in laboratory settings where collecting data is relatively straight-forward. However, using consumer sensors in naturalistic settings that present significant challenges, including noise artefacts and missing data, has not been as extensively investigated. Additionally, the majority of prior studies focused on wrist-worn optical HR sensors. Arm-worn sensors have not been extensively investigated either. In the present study, we validate HR measurements obtained with an arm-worn optical sensor (Polar OH1) against those obtained with a chest-strap electrical sensor (Polar H10) from 16 participants over a 2-week study period in naturalistic settings. We also investigated the impact of physical activity measured with 3-D accelerometers embedded in the H10 chest strap and OH1 armband sensors on the agreement between the two sensors. Overall, we find that the arm-worn optical Polar OH1 sensor provides a good estimate of HR (Pearson r = 0.90, p <0.01). Filtering the signal that corresponds to physical activity further improves the HR estimates but only slightly (Pearson r = 0.91, p <0.01). Based on these preliminary findings, we conclude that the arm-worn Polar OH1 sensor provides usable HR measurements in daily living conditions, with some caveats discussed in the paper.

## 1. Introduction

Consumer wearable sensors can help people monitor their overall health and provide valuable information for prevention of severe diseases and injuries.^1–3^ Cardiovascular parameters such as heart rate (HR) and heart rate variability (HRV) are among the most common physiological measures that people track with their wearable sensors. The HR and HRV measurements captured by the sensors not only provide information about physical health, they can also help track mental stress which has secondary deleterious effects on health, including mental health.^4–7^ The most commonly used sensors for cardiovascular measurements are wrist-worn smart watches; however, chest strap sensors are also widely used, especially in the context of sports. More recently, a class of devices that are designed to be worn on the upper arm or the forearm have become commercially available. The wrist-worn and arm-worn sensors rely on photoplethysmography (PPG: optical sensing) and chest strap sensors rely on electrocardiography (ECG). The latter tend to be more accurate than the former.^3,4,8–11^ While there has been extensive prior work validating wrist-worn heart rate sensors, most of this work has been done in laboratory conditions.^1,4,12^ Less work has been done to examine the validity of optical HR sensors in completely unconstrained and uncontrolled naturalistic settings. For example, a recent meta-review of 44 studies that reported on validity of wrist-worn optical sensors found only 7 studies that included daily living activities outside of a lab setting.^13^ Furthermore, the results of this work have been mixed with respect to the ability of optical sensors to accurately measure heart rate in these unconstrained conditions.^2,14–17^ Even fewer studies have examined arm-worn devices as an alternative to wrist-worn sensors.^18–21^ These studies focused mainly on the use of these devices in the context of sports activities and demonstrated that armband devices are robust to even very strenuous physical activity. For this reason, we selected the Polar OH1 armband as an alternative to wrist-worn devices. Our current study aims to add to this prior literature a preliminary investigation of an armband optical heart activity sensor worn for an extended period of time in everyday life settings. We use the Polar OH1 armband sensor together with the Polar H10 chest strap sensor as a reference device to collect PPG, ECG and accelerometer data and explore the feasibility and accuracy of using Polar OH1 armband’s PPG measurements obtained in the naturalistic environment with Polar H10 chest strap’s ECG measurements as the reference standard. Additionally, we aimed to examine the impact of motion on the accuracy of HR estimates.

## 2. Study Design

This preliminary study is part of a larger study of cigarette smokers. The larger study is ongoing and is aimed at predicting smoking events in order to develop or use therapeutic interventions (e.g., nicotine lozenge) that can be administered just-in-time. In this study, participants are asked to wear several sensors including the Polar H10 and OH1 for approximately 14 consecutive days (2 weeks). During both weeks, the participants are asked to use a smartphone app specifically designed for this study (PhysiAware^®^) that uses Bluetooth Low Energy (BLE) interface to connect to the study devices, collect the real-time measurements and transmit them to a study server several times a day. The participants are also asked to use the app to indicate when they smoke each cigarette and the reasons for smoking the cigarette. The current analysis includes only Polar H10 and OH1 heart activity trace and accelerometer data from the first 16 participants from the larger study.

## 3. Data Collection and Processing

This study was approved by the University of Minnesota Institutional Review Board and is currently ongoing.

### 3.1. Data Collection

The data collected from the armband and chest strap sensors are temporarily stored by the PhysiAware^®^ app on the smartphone of each participant until the participants upload the data to a University of Minnesota server. The PhysiAware^®^ app was developed specifically for this project as a native app on iOS and Android platforms. Figure 1 illustrates both versions and shows the main screen that the participants would see after logging in with their study credentials (a randomly generated study ID).

**Fig. 1:**
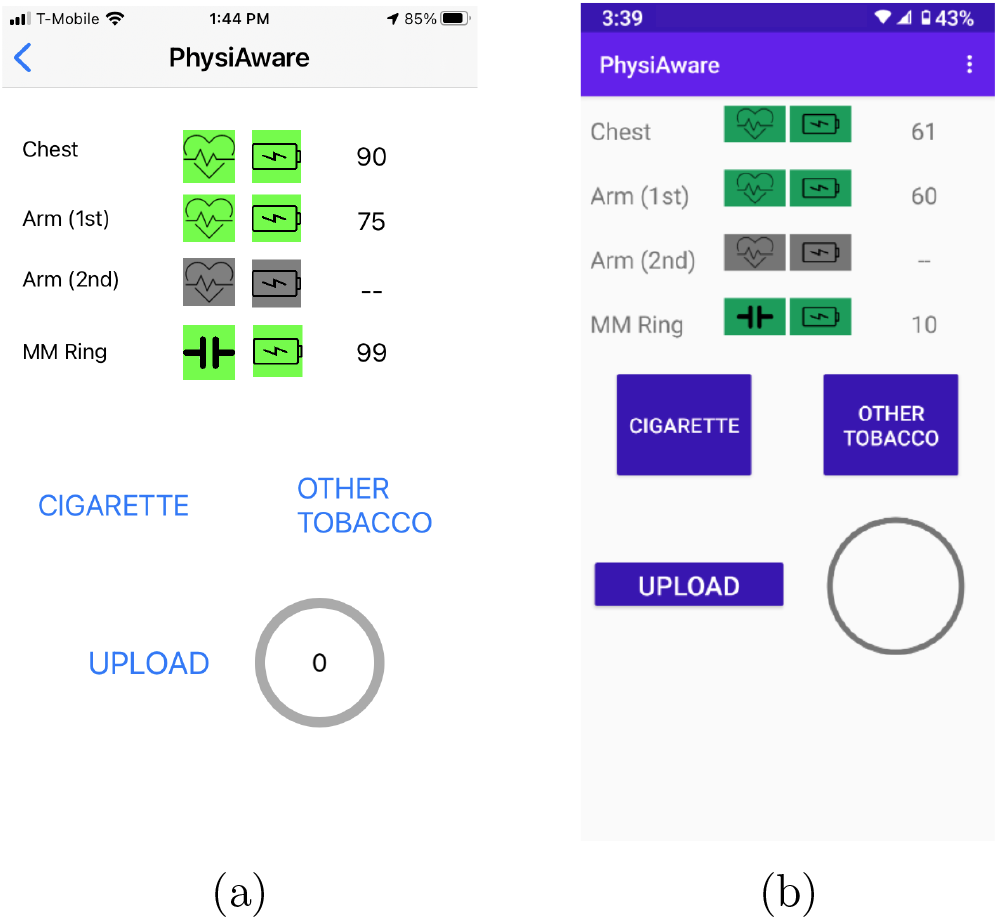
The iOS and Android versions of the PhysiAware app. a) iOS. b) Android. ^*∗*^ MM Ring is a ring sensor (MoodMetric^®^) used in the study for monitoring electrodermal activity - not relevant to the current analysis.

The iOS and Android apps implemented the standard Polar BLE application programming interface for collecting raw data from the sensors. Both the H10 chest strap and the OH1 armband sensors are capable of storing limited data in their onboard memory; however, this capability is not open to third-party developers when the OH1 armband sensor is operated in the “PPI” mode (i.e., the mode that calculates and reports inter-beat interval durations needed for heart rate variability measurements). For our study, we wanted to leverage the “PPI” mode specifically along with collecting raw blood volume pulse data. Thus, all sensor data were streamed “live” over the BLE interface rather than stored locally and transferred in batches. The streaming data were aggregated on the smartphone and the participants were prompted every 3-4 hours to upload their data to the study server. The motivation for not doing automated uploads stems from the fact that some of the participants may have limited or costly cellular data plans. Therefore, we designed the app to detect the presence of Wi-Fi connectivity and alert the participants only when a Wi-Fi network (vs. a cellular data network) was available to upload the data. We also wanted to provide the participants with the ability to manually control the uploads as they take a significant amount of bandwidth and can be disruptive to the participant’s other activities on their smartphone. These design considerations were adopted in order to make the app as accessible as possible to a broad range of participants from a variety of socioeconomic backgrounds.

Due to the remote and naturalistic nature of the study, we encountered several other challenging issues that affected data collection. For example, Polar OH1 arm sensor battery life is approximately only 8 hours, which precludes continuous monitoring. To compensate for this challenge, each participant was provided with two OH1 sensors that they could use interchangeably while the other sensor was being charged. While all participants participated in a remote training session via Zoom with the study coordinator on how to wear and use the study devices, situations arose where participants unintentionally corrupted the data. This included wearing the sensors incorrectly, or forgetting to wear them at all. These challenges are inherent to remote naturalistic settings and result in lower volume of usable data than what can be obtained in laboratory conditions or with extensive hands-on training. The collected data still may present further challenges due to noise from the variability of the environments and participants’ daily life activities.^22^

### 3.2. Data Processing

For the current study, we selected the signals available simultaneously from both the H10 chest strap and OH1 armband to time-align the two signals as illustrated in Figure 2. Some of the more frequent noise artifacts included short gaps with missing samples. The majority of these gaps were under 60 seconds in duration and were likely attributable to interruptions in BLE connectivity between the smartphone and the sensors. The missing data corresponding to these short gaps that are under 60 seconds comprises on average 0.52% of the total data volume for ECG and 0.94% for PPG. Our current approach for dealing with these short gaps is to back- and forward-fill them by taking half of the values needed to fill the gap from the preceding signal, and the other half from the subsequent signal. This approach is motivated by the thought that the HR signal does not typically change dramatically over a short period of time; however, if such a change does occur during the 60 second gap (e.g. the participant begins strenuous activity during the gap) by forward-filling the first half of the gap and back-filling the second half we expect to represent the start time of the increase in HR more accurately. Less frequent gaps longer than 60 seconds were left as missing data and excluded from analysis.

We collected the high frequency time indexed ECG data and 3-D acceleration data from Polar H10 chest strap sensor at a sampling rate of 130Hz and 200Hz, respectively. The time indexed PPG and 3-D acceleration data from the Polar OH1 armband were sampled at 135Hz and 50Hz, respectively. To detect peaks in the ECG and PPG signals and calculate instantaneous HR we used Kubios^23^ software package (version 3.5.0) which also performs additional noise filtering and generates HR estimates in the output. Other computational approaches were considered such as band pass filtering, detrending methods for removing noise, and manually calculating HR with code. However, we opted to use the Kubios for preprocessing so that our results are more easily reproducible and applicable to a wider research audience. Kubios processes ECG and PPG data by using an automatic beat detection algorithm and HR calculation from inter-beat intervals. Additionally, Kubios applied a detrending approach on the ECG and PPG data based on smoothness priors regularization. The detrending method removes the slow non-stationary component of the signals.^24^ Peak detection and time-alignment between PPG and ECG signals are shown in Figure 2. Less frequent gaps longer than 60s were filled with zeros prior to Kubios processing to maintain the time-alignment between the ECG and PPG data. Kubios generates ‘NaN’ values for heart rate variability features for the zero-filled sections resulting in excluding these sections from analysis.

**Fig. 2:**
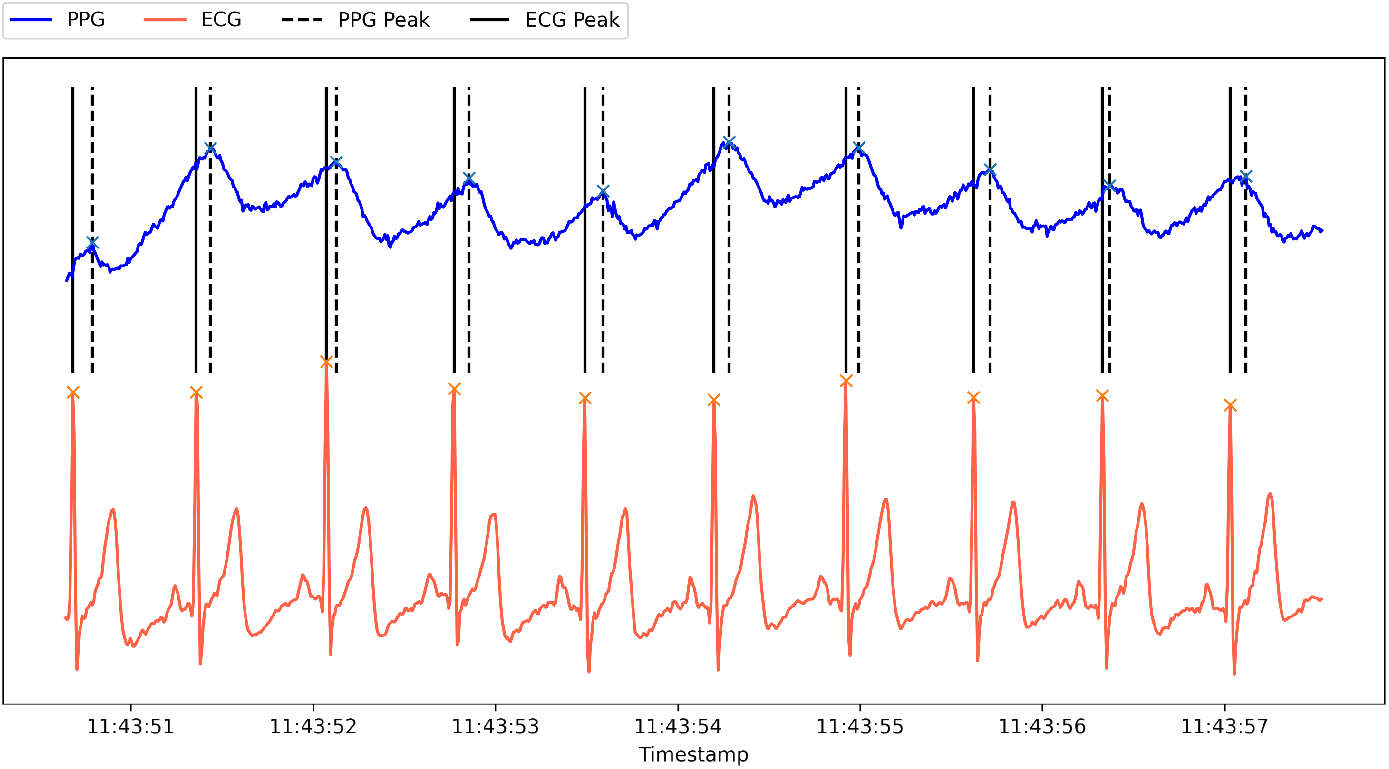
Illustration of peak alignment between ECG (lower) and PPG (upper) signals.

After handling missing data as described above, we imported the data into Kubios to generate HR estimates over 1-minute frames overlapping by 10 seconds. The resulting time series were used to calculate the Pearson correlation coefficients between ECG and PPG HR estimates. Due to the large size of the high frequency time series ECG and PPG data (e.g., up to 120 million rows for 11 days of ECG data per participant), we segmented the data into smaller chunks ranging from a few hours to one day (24 hours) before importing into Kubios.

### 3.3. Filtering

To investigate the impact of noise introduced by physical activity in daily life, we experimented with several physical activity filters based on 3-D accelerometer data from accelerometers embedded in the H10 chest-strap and OH1 armband devices. Since the H10 is attached to the person’s torso and OH1 is attached to the upper forearm, we expect that these sensors will capture different and potentially complementary types of activity. All physical activity filters use the magnitude of acceleration along the x, y, z axes from a given 3-D accelerometer. The calculation of the magnitude of overall acceleration is as follows:

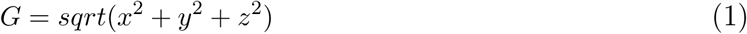

where G represents the magnitude of acceleration and x, y, z represents acceleration along each of the 3 axes. In the rest of the paper we refer to this overall magnitude of acceleration as G-value. This measure is used to filter out the data that contain high levels of physical activity defined as above the 75th percentile of all G-values for a given sensor. We then defined three filters based on the G-values calculated from a) H10 chest strap, b) OH1 armband, and c) from the union of both H10 and OH1 G-values (i.e., when either the chest-strap or the armband device indicated excessive motion). Each of the three filters also removes samples that result in HR greater than 200 beats per minute (based on maximum heart rate calculated as 211 - (0.64*age)^25^) or lower than the empirically determined 3rd percentile. Activity filters were used in this study to assess if the correlation between the measures reported by the PPG and ECG sensors are adversely impacted by including periods of physical activity so as to assess the quality of the data collected by the arm-band sensor during times of physical activity.

## 4. Results

The basic demographic and physiological characteristics of participants are presented in Table 1. The participant characteristics presented in Table 1 demonstrate the variability present in HR measurements across study participants.

**Table 1:** Participant Characteristics

| ID | Age | Sex | Race | Arm | HR (OH1) | HR (H10) | G-val (OH1) | G-val (H10) |
| --- | --- | --- | --- | --- | --- | --- | --- | --- |
|  |  |  |  |  | Mean (SD) | Mean (SD) | Mean (SD) | Mean (SD) |
| 1 | <40 | m | non-white | left | 83.56 (10.0) | 85.29 (11.0) | 1007 (23) | 973 (14) |
| 2 | <40 | f | white | left | 76.28 (9.0) | 77.11 (9.0) | 1021 (12) | 1007 (17) |
| 3 | 40-50 | f | white | left | 77.43 (6.0) | 77.27 (6.0) | 968 (137) | 1015 (7) |
| 4 | <40 | m | non-white | left | 69.17 (8.0) | 69.92 (9.0) | 999 (15) | 994 (11) |
| 5 | 40-50 | f | white | left | 96.73 (7.0) | 96.84 (7.0) | 1002 (9) | 988 (29) |
| 6 | >50 | f | white | left | 74.55 (7.0) | 75.38 (7.0) | 1001 (12) | 995 (13) |
| 7 | <40 | m | white | left | 106.9 (8.0) | 107.88 (8.0) | 1023 (11) | 999 (28) |
| 8 | <40 | f | white | left | 103.45 (5.0) | 103.85 (5.0) | 1012 (13) | 993 (10) |
| 9 | <40 | m | white | left | 94.02 (10.0) | 96.14 (10.0) | 1002 (18) | 1019 (11) |
| 10 | 40-50 | f | white | left | 79.88 (9.0) | 80.73 (9.0) | 1017 (6) | 979 (17) |
| 11 | <40 | m | white | left | 94.22 (14.0) | 94.85 (14.0) | 1024 (22) | 1019 (11) |
| 12 | 40-50 | m | white | left | 84.0 (8.0) | 85.09 (8.0) | 1008 (12) | 1006 (12) |
| 13 | >50 | m | white | right | 86.65 (8.0) | 88.02 (9.0) | 1019 (66) | 1009 (10) |
| 14 | <40 | f | white | left | 78.8 (9.0) | 79.71 (9.0) | 1024 (14) | 976 (16) |
| 15 | <40 | f | white | left | 76.19 (9.0) | 76.75 (9.0) | 1017 (16) | 997 (20) |
| 16 | 40-50 | f | white | left | 96.56 (6.0) | 97.02 (6.0) | 1013 (8) | 991 (10) |
| **Mean** | 42 | – | – | – | 86.14 (8.3) | 87.0 (8.5) | 1009.8(25) | 997.5 (15) |

### 4.1. Agreement between PPG and ECG HR Estimates

We used the HR values generated by Kubios for both PPG and ECG signals and calculated Pearson correlation of the two sets of HR values. The Pearson correlation measures the strength of the linear relationship between two variables. As shown in Table 2, the HR calculated from PPG data is positively correlated with the HR calculated from ECG data with correlation coefficients higher than 0.80 except for one participant’s data likely due to excessive motion or the chest strap not being tight enough. The results summarized in Table 2 show that the majority of participants’ correlation coefficients increased only modestly after using physical activity filters. All correlations in Table 2 are statistically significant (p-value *<*0.01).

**Table 2:** Correlation between ECG and PPG estimates of HR and number of samples remaining after applying various physical activity filters based on accelerometers embedded in devices.

| ID | Physical Activity Filters |  |  |  |  |  |  |  |
| --- | --- | --- | --- | --- | --- | --- | --- | --- |
|  | No filter |  | OH1 armband |  | H10 chest strap |  | OH1+H10 |  |
|  | Corr. | N* | Corr. | N | Corr. | N | Corr. | N |
| 1 | 0.82 | 4021 | 0.82 | 3146 | 0.85 | 3274 | 0.86 | 2600 |
| 2 | 0.95 | 2195 | 0.95 | 1950 | 0.94 | 1649 | 0.94 | 1466 |
| 3 | 0.97 | 124 | 0.97 | 101 | 0.98 | 92 | 0.98 | 79 |
| 4 | 0.95 | 280 | 0.94 | 216 | 0.98 | 245 | 0.98 | 198 |
| 5 | 0.96 | 326 | 0.96 | 291 | 0.96 | 275 | 0.96 | 252 |
| 6 | 0.83 | 2025 | 0.84 | 1762 | 0.85 | 1768 | 0.86 | 1544 |
| 7 | 0.95 | 64 | 0.96 | 60 | 0.77 | 44 | 0.76 | 42 |
| 8 | 0.93 | 177 | 0.93 | 148 | 0.92 | 139 | 0.92 | 120 |
| 9 | 0.53 | 2312 | 0.55 | 2026 | 0.58 | 1506 | 0.58 | 1298 |
| 10 | 0.98 | 26 | 0.98 | 25 | 0.97 | 25 | 0.98 | 24 |
| 11 | 0.96 | 114 | 0.99 | 80 | 0.99 | 76 | 0.99 | 63 |
| 12 | 0.93 | 3219 | 0.93 | 2395 | 0.92 | 2813 | 0.92 | 2102 |
| 13 | 0.92 | 569 | 0.94 | 536 | 0.95 | 292 | 0.95 | 269 |
| 14 | 0.96 | 613 | 0.97 | 514 | 0.96 | 559 | 0.97 | 479 |
| 15 | 0.93 | 677 | 0.93 | 615 | 0.95 | 530 | 0.95 | 489 |
| 16 | 0.90 | 1455 | 0.93 | 1264 | 0.90 | 1378 | 0.94 | 1196 |
| **Unweighted Mean** | 0.90 | 1137.3 | 0.91 | 945.5 | 0.90 | 916.6 | 0.91 | 763.8 |
| **Weighted Mean** | 0.85 | 1137.3 | 0.85 | 945.5 | 0.87 | 916.6 | 0.87 | 763.8 |

\* number of samples (HR estimates) used to calculate the correlations. Each sample corresponds to HR calculated over a 1-minute frame and, thus, N also approximates the number of minutes of data included in the correlation analysis, not counting the overlaps between frames.

The average HR correlation before any physical activity filtering is 0.90. Using either or both the OH1 armband or H10 chest strap physical activity filter to remove data with high G-values increases the average HR correlation to 0.91. Since different participants have different numbers of samples, we also report correlations weighted by the number of samples resulting in slightly lower estimates but still remaining above 0.80 (see Table 2).

Correlations between PPG and ECG heart rate estimates by age and sex of the participants are illustrated in Figure 4 and suggest that these participant characteristics did not substantially affect the results.

**Fig. 3:**
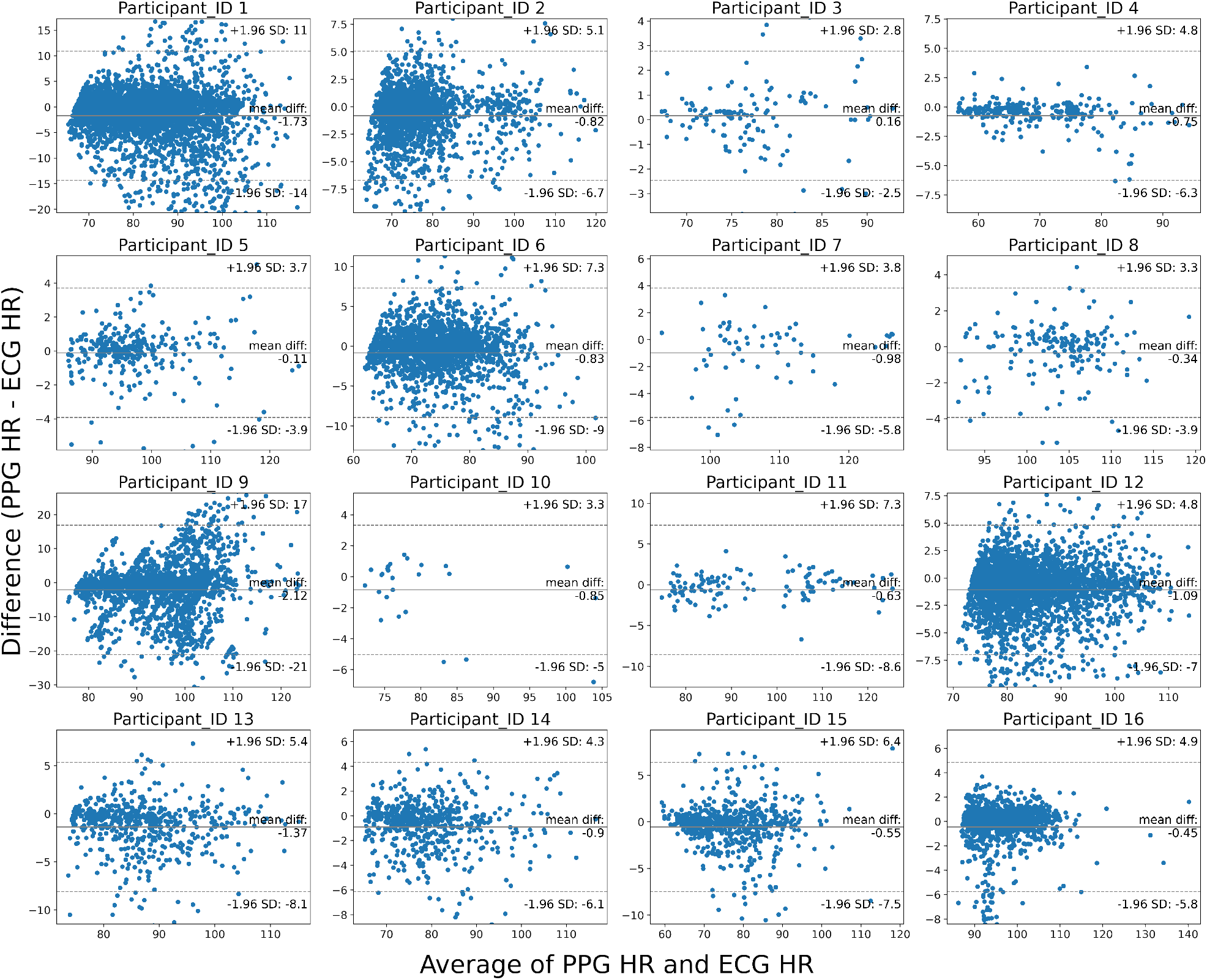
Bland-Altman Plot of the differences in PPG and ECG estimates of HR without any filtering physical activity.

**Fig. 4:**
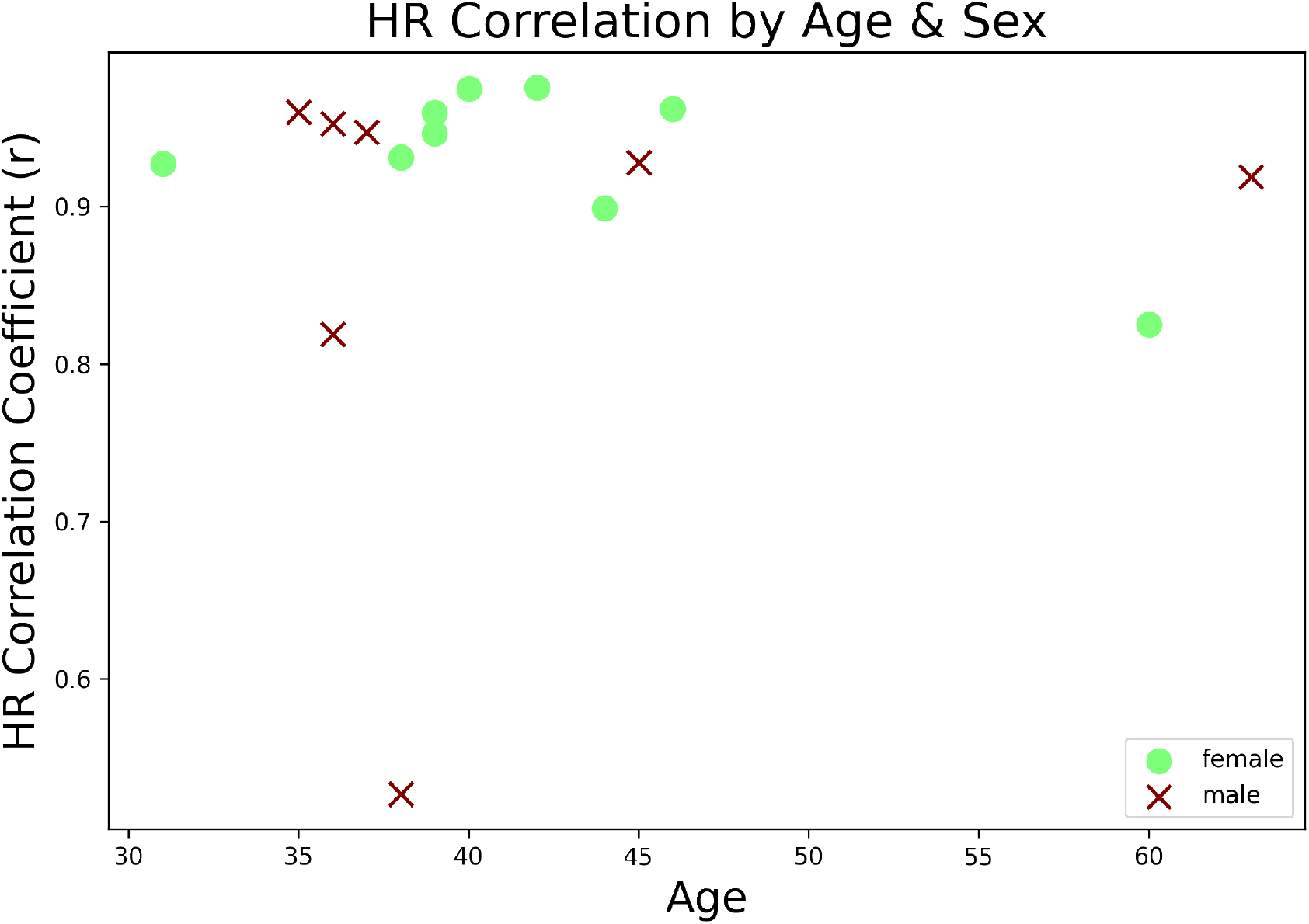
Differences in correlations between PPG and ECG HR by age & sex of the participants.

## 5. Discussion

The results of our preliminary study indicate that it is feasible to obtain HR estimates from the Polar OH1 armband that are in agreement with reference ECG estimates. These findings are encouraging for studies that involve observation of cardiac activity in naturalistic settings.

Several previous studies^26–28^ that examined the agreement between PPG and ECG signals reported correlations between 0.91 and 0.98 which is comparable to the correlations we found in the present study; however, some previous studies noted somewhat variable performance of wrist-worn PPG sensors with some devices having correlation with ECG in the 0.83-0.84 range^16^ or even lower.^17^ Furthermore, while the ecological study by Nelson et al.^26^ demonstrated overall low error rates for wrist-worn (Apple Watch 3 and Fitbit Charge 2) devices under sleeping, sitting, walking, and running conditions over a 24 hour period, they also found relatively high error rates during activities of daily living and as movement became more erratic during various conditions. These decreases in accuracy are likely due to the fact that wrist-worn devices by design tend to fit relatively loosely around the wrist. An overly tight fit would make the device uncomfortable to wear for long periods of time. The Polar OH1 armband used in our study seeks to overcome this problem by placing the sensor higher up the forearm which tends to experience lower amplitude motion than the wrist during activity and allows for tighter fit with an elastic strap that also minimizes the amount of motion. Another key difference in our study is the length of the observation period. In our study, participants wore the sensors as continuously as they were comfortable with over an approximately two-week period. We are not aware of other studies in which participants wore a chest strap and an armband for such an extended period of time. The extended nature of the observation period enables us to examine the performance of the armband sensor in a greater variety of naturalistic conditions and activities of daily living. The fact that we find the armband to provide accuracy on par with other devices used in laboratory conditions is particularly encouraging as they show that this type of PPG sensor can be comfortably worn over a long period of time and provides reliable measurements.

We also find some individual variation across participants as illustrated by the correlations reported in Table 2. Additionally, the Bland-Altman plots in Figure 3 show there’s also some variability in the distribution of the differences between ECG and PPG HR estimates across the participants. However, the mean differences of all participants are close to zero and majority of the data is within the 95% confidence intervals of limits of agreement. The distribution of the points outside of the 95% confidence intervals does not suggest a clear pattern of association between the differences and the magnitude of the heart rate measurements.

Visual examination of the differences between groups by age and sex, shown in Figure 4, suggests no substantial differences between these groups. Due to the small number of participants, we did not perform a formal statistical subgroup analysis. The data shown in Figure 4 suggests that the male subgroup contains a possible outlier (participant 9 in Table 2).

Several prior studies examined the impact of skin tone on optical green wavelength heart rate sensor accuracy and found that these sensors were reasonably accurate across various skin tones but slightly less accurate for darker skin tones varying by devices and conditions.^13,29,30^ While our study so far included only 2 participants with non-white skin tone, our results are consistent with this prior work in that we found that the optical OH1 armband was only slightly less in agreement with the ECG estimates of heart rate than the group average for one of the two non-white participants (r = 0.82 vs r = 0.90 - see Table 2, participant 1 in “No filter” column) and slightly higher than the group average for the other non-white participant (r = 0.95 vs r = 0.90 - see Table 2, participant 4 in “No filter” column). Clearly, we cannot draw any definitive conclusions from these results due to the small number of participants overall and non-white participants in particular.

Another important finding is that removing heart data that corresponds to physical activity based on the accelerometer values did not have a major impact on the agreement between ECG and PPG estimates of heart rate. This is an encouraging finding because it indicates that the armband sensor provides robust heart rate estimates in the presence of physical activity.

## 6. Limitations and Challenges

The results should be interpreted in light of several limitations. First, our sample size is small as this is a preliminary pilot study. Second, we use only the standard accelerometer-based filtering techniques. More advanced filtering techniques exist that may be able to further reduce noise, potentially increasing the correlation. Finally, the participants included in this study are smokers, whose physiological characteristics may differ from the general population.

In addition to the limitations listed above, we also want to highlight several valuable lessons that we learned in the process of doing this study that can be applied in our future work or by others who intend to perform similar studies. On the technical side, we found that some of the participants had some trouble with maintaining the connectivity between the wearable sensors and the smartphone. The Bluetooth devices used in this study have a relatively short range (10-30 meters); therefore, the “live” streaming mode for data transfer is vulnerable to the participants walking away from their smartphones beyond the Bluetooth range. Clearly, this results in undesirable data loss which may be prevented by recording data locally on the sensor devices in addition to streaming.

We also found significant differences in terms of the technical challenges in app development for the two platforms: iOS and Android. While developing for the Android platform was logistically easier than for iOS mostly due to complicated security controls on iOS apps, it was also much more challenging to use Android apps in the study due to large variability in how various Android smartphone manufacturers handle battery management. In order to maintain the streaming of data from devices to the smartphones, the battery management mode had to be manually turned off by the participants and was achieved with variable success depending on which smartphone the participant owned. This issue is more difficult to resolve without resorting to recruiting only participants who own Apple smartphones, which would make recruitment more difficult and may introduce unintended selection bias into the study. In the current study, we addressed this issue by monitoring incoming data on a regular basis for signs of significant data loss and had the study coordinator follow up with those participants that were identified this way.

## 7. Acknowledgments

Funded by NIH NIDA award DA049446 and supported by the University of Minnesota CTSI (UL1-TR002494). The content of this article is solely the responsibility of the authors and does not necessarily represent the official views of the NIH.

## Notes

### Competing Interest Statement

The authors have declared no competing interest.

